# Dispersal-induced persistence of chaotic ecosystems under extreme climatic events

**DOI:** 10.1101/2025.07.29.667107

**Authors:** Dweepabiswa Bagchi

## Abstract

Ecosystems are strongly disrupted by large-scale natural disturbances called extreme events which can potentially lead to their local extinction. In this work, we study one of the most important examples of this, namely, a coastal ecosystem interacting with a tropical cyclone, one of the strongest extreme weather events. We show, that the ecosystem, while perfectly self-sustainable when unperturbed, nevertheless undergoes a reduction in habitat quality when impacted by the extreme event. We show that a certain degree of dispersal between the affected patch and a less affected patch leads to non-zero population density, whereas the population of an uncoupled but affected patch suffers extinction. We also demonstrate that dispersal between two equally affected coastal ecosystems, eventually leads to the extinction of their populations. We show that the critical degree of dispersal for the persistence of the ecosystems under the onslaught of extreme events increases with both damage-susceptibility of the ecosystem and ferocity of the extreme event. We also analyze the dynamical behavior of the affected coastal ecosystem under increasing dispersal.

## I. INTRODUCTION

The dispersal of species among the patches of an ecosystem creates an ecological network leading to a metacommunity. Intriguingly, dispersal is one of the major factors determining the population dynamics of the entire network [1]. Generally, dispersal has a stabilizing influence on the overall metapopulation dynamics [2], which leads to a decreased risk of extinction [3, 4]. Numerous studies have also shown that large disruptions in the population dynamics of any species of the metacommunity can drastically decrease the population density that is significant enough to drive the species to extinction [5–12]. This is devastating to the persistence of not only the species itself, but the whole ecosystem. Clearly, the dynamical impacts of such disruptions and those of dispersal contradict each other. Therefore, from an eco-conservation point of view, the role of dispersal in countering the critical decrease in the population density caused by large-scale disruptions merits analysis and investigation.

Extreme events are characterized by very irregular and erratic behavior of observables of a system. The observable in question takes on an ‘extreme’ i.e., unrealistically high (or low) values during the event. From an ecologically relevant perspective, large-scale disturbances are classified as extreme events [13]. The most common among them to strongly affect ecosystems are floods, tsunamis, cyclones, forest fires, and epidemics [14]. A plethora of studies have demonstrated that such events can cause large enough disruptions that can critically lower the population density of the species and consequently drive the corresponding ecosystem to extinction [8–10, 15–19]. Recent studies have also shown that the onslaught of such extreme events is rapidly increasing in frequency [20, 21]. Two major ecologically relevant extreme events are tropical cyclones and forest fires [22, 23]. Motivated by this, we investigate the impact of a tropical cyclone and the resulting storm surge on a coastal ecosystem [24, 25], and the impact of habitat destruction due to forest fires on a terrestrial ecosystem [26, 27].

Notably, as far as impacts on ecosystems are concerned, detrimental extreme events also include manmade disasters. For a (benthic) coastal ecosystem, one of the most important extreme events is the spill of chemicals, for instance, the spill of crude oil [28]. The subsequent approach one can undertake will be analyzing the role of dispersal in reviving ecosystems with decreasing habitat quality using an end-to-end model to the case of chemical spills in coastal ecosystems [29]. The aftereffects of such sudden decrease in habitat quality are detrimentally long term and large-scale [30, 31]. Studies have shown that habitat quality is also dangerously degraded in such cases [32].

Even though large dynamical disruptions caused by extreme events can lead to the extinction of a species, in reality, a considerable number of ecosystems persist in spite of being impacted regularly and strongly by extreme events [33]. Two approaches are the most prevalent in existing modeling studies that intend to understand the underlying reason for the survival of such ecosystems. Some studies focus primarily on ecological dynamics and introduce an ad hoc, simple dynamical variable representing the environment [34],

[35] into an ecosystem model. Such models, however, are often simplistic and sometimes even one-dimensional. The absence of the complexity associated with climatic or environmental dynamics in these models, makes them inappropriate to meaningfully represent complex phenomena like extreme events. A few existing studies which consider both extreme events and ecosystem models in some detail also examined only the immediate observable changes in the ecosystem [9], [36]. Some studies considered only species-specific impacts of extreme events despite the species in question being a part of a complex food web [37– However, to the best of our knowledge, so far there have been no studies that simultaneously consider bio-diverse ecosystem models, and nonlinear complex models of extreme events that strongly impact the former along with ecological mechanisms that ensure sustained survival of such ecosystems. A model incorporating complexities from all ends of a natural phenomenon is called an ‘end – to - end model’ [40], [41]. Our study deals with such an end-to-end models corresponding to one typical extreme event.

In this study, we consider a coastal ecosystem exhibiting chaotic dynamics when unperturbed, but changes its course of evolution when detrimentally impacted by a tropical cyclone. At first, we examine the effect of dispersal between two identical affected coastal ecosystems. Then, we investigate the effect of dispersal between a particular affected coastal ecosystem and safer inland ecosystems by estimating the critical dispersal required to retain the persistence of the affected ecosystems. We find that the affected patches survive from the extinction above a critical dispersal rate, which increases with the increase of both intensity of the extreme event and damage-proneness of the ecosystem. We also investigate the effect of increasing dispersal between a coastal ecosystem and inland ecosystem. The plan of the paper is as follows. In Sec. II, we introduce a cyclone boundary-field model along with a storm-surge model that affects a coastal ecosystem. Both the affected coastal ecosystem and the inland ecosystems are described by a modified Powell-Hastings model [42, 43]. We discuss our results in Sec. III and provide discussion and conclusions in Sec. IV.

## II. MODELS

### A. Tropical cyclone impact on an ecosystem

#### 1. Cyclone boundary-field model

Tropical cyclones are essentially huge volumes of spiraling wind that rotate about a zone of low pressure. The boundary wind field of a tropical cyclone is described by a modified Navier-Stokes model [24, 44].The time evolution of the wind-field of a tropical cyclone (*u, v*) is governed by the equations of motion

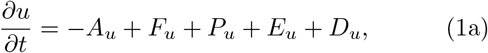

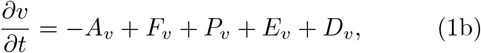

where, 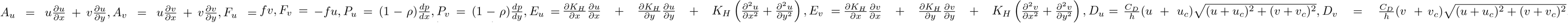. The pressure gradient terms (*P*_*u*_, *P*_*v*_) are calculated by transforming the radial derivative of the Holland pressure [45] into the Cartesian coordinates (*x, y*). The radial pressure derivative term is given as,

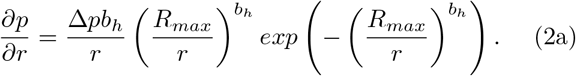

. The dynamical equations include the advection terms, *A*_*u*_ and *A*_*v*_, accelerations induced by the Coriolis forces *F*_*u*_ and *F*_*v*_, pressure gradient terms *P*_*u*_ and *P*_*v*_, fluid viscosity terms *E*_*u*_ and *E*_*v*_ and surface-drag coefficient terms *D*_*u*_ and *D*_*v*_. Barring specific cases, the ‘Holland parameter’, *b*_*h*_ is fixed as *b*_*h*_ = 2.1 and the pressure deficit is fixed as Δ*p* = 4350. The other parameter values are fixed as *f* = 0.06, *ρ* = 1.15, *C*_*D*_ = 0.05, *h* = 1000 throughout the manuscript unless otherwise specified. The Horizontal eddy viscosity coefficient *K*_*H*_ negligibly affects the wind field model [24] and so, it can be assumed to be a constant. We consider *K*_*H*_ = 0.04.

The wind-field (*u, v*) from the cyclone creates a huge atmospheric pressure called the wind-stress, defined as 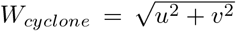. The wind pressure (denoted by *W*_*cyclone*_), creates an upwelling of water called storm surge, modeled by the El-Nino Southern Oscillation (ENSO) model elaborated in the following section. A parameter defined as *w*_*cyclone*_(∝ *W*_*cyclone*_) is included in the ENSO model to quantify the storm surge.

#### 2. Storm-surge model

When a cyclone forms over an ocean, the huge wind-stress facilitates the elevation of ocean water levels, called storm surge, which can be modeled by the El-Nino Southern Oscillation model [25]. The wind-field (*u, v*) from the cyclone creates a huge atmospheric pressure called the wind-stress, defined as 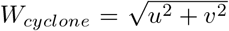. When the cyclone forms over ocean, this huge stress facilitates the elevation of ocean water levels. This elevated ocean water level is called the storm surge, which is modeled by the following El-Nino Southern Oscillation (ENSO) model [25]

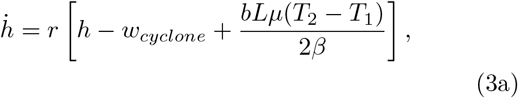

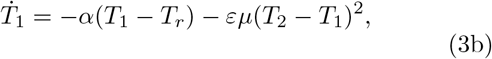

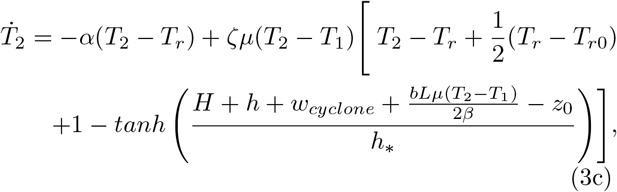

where, *h* corresponds to the height of the thermocline water level of the ocean, and *T*_1_ and *T*_2_ are the equatorial ocean surface temperatures [25]. The parameter values are chosen as: *r* = 1, *β* = 200, *α* = 1, *ε* = 0.0961, *µ* = 0.0026, *H* = 100, *z*_0_ = 75, *h*_*_ = 62. In Eq. (3c), the term *w*_*cyclone*_ is proportional to the cyclonic wind-stress *W*_*cyclone*_ by a factor of 0.001. The height *h* of the up-welling water, storm surge, affects the habitat quality of the impacted coastal ecosystem. In this work, the carrying capacity *K*_0_ quantifies the degree of damage to the habitat quality of the coastal ecosystem.

#### 3. Model of metacommunity impacted by tropical cyclones

A coastal (or benthic) ecosystem is the primary frontier for the storm surge impact. The food web of both the coastal and inland ecosystem is described by a modified Powell-Hastings model [42, 43].

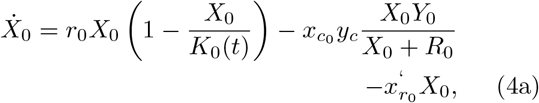

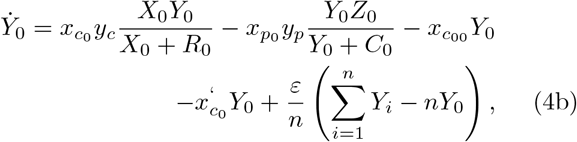

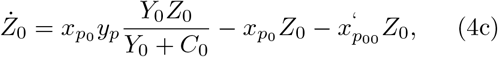

where *X*_0_, *Y*_0_, *Z*_0_ are the prey, predator and consumer of the food web, respectively. *r*_0_ is the growth rate of prey. Among the other parameters, 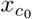 is the mortality rate of the prey *X*_0_, *y*_*c*_ is the predation rate of the predator *Y*_0_ on 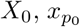 is the mortality rate of the predator *Y*_0_, *y*_*p*_ is the predation rate of the consumer *Z*_0_ on *Y*_0_, *R*_0_ and *C*_0_ are the half saturation constants of *X*_0_ and *Y*_0_, respectively. The number of habitats is denoted by *n*. Throughout this work, unless otherwise specified, the parameter values are fixed as 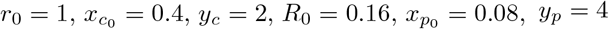.

Other than the mortality terms in the model, we further introduce a gradually diminishing mortality term, induced by increase in the salinity and toxicity by the storm surge for prey, predator and consumer. The storm surge induced mortality rate of the prey is chosen as 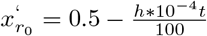. The corresponding terms for predator and consumer is chosen as 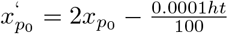 and 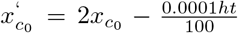, respectively. It is reasonable to consider that the extra predator and consumer mortality terms are proportional to the individual’s inherent mortality rate. The dispersal term 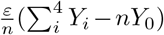 quantifies the dispersal between the coastal ecosystem and the inland ecosystems, where *Y*_*i*_ corresponds to the predator density of the latter. It is, however, to be noted that we also consider a case where one affected coastal ecosystem is connected with another, in that case the dispersal term is given by 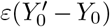. The orientation of all the patches is given by Fig. 2.

**FIG. 1.**
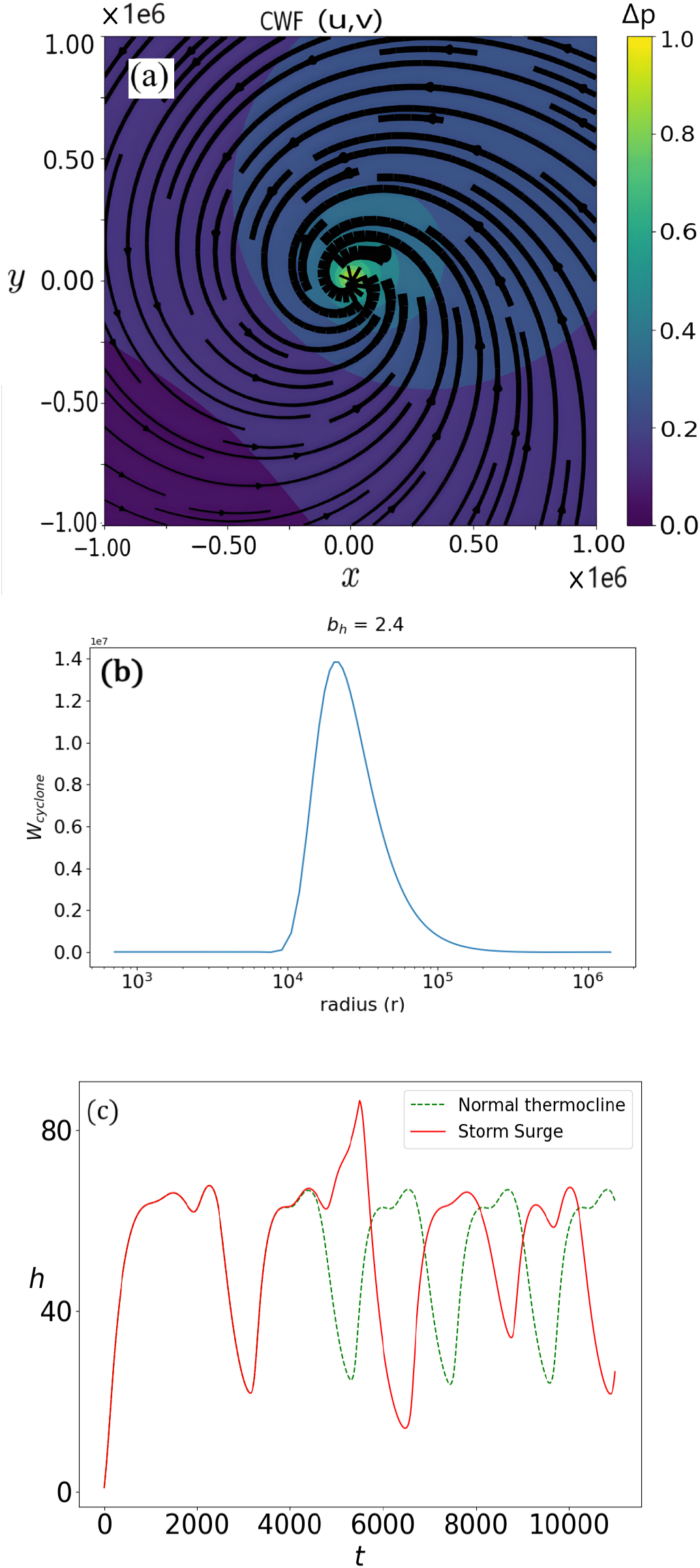
(a) The cyclonic wind field in (*x, y*) plane. The black arrows show the direction of the wind, thickness of the arrows signify relative intensity/speed and the colored contour plot shows the pressure deficit that generates the cyclone. (b) Radial wind-stress profile of an intense tropical cyclone (Holland parameter: *b*_*h*_ = 2.4). The *r*-axis is in *log* scale. The parameter values are the same as in Eqs. (1)-(2). (c) (green line) Periodic variation of thermocline ocean water level without cyclonic wind-stress. (red line) Storm surge in thermocline height formed by wind-stress from a cyclone with the Holland parameter *b*_*h*_ = 2.1. All the other parameters are the same as in Eqs. (1)-(3).

**FIG. 2.**
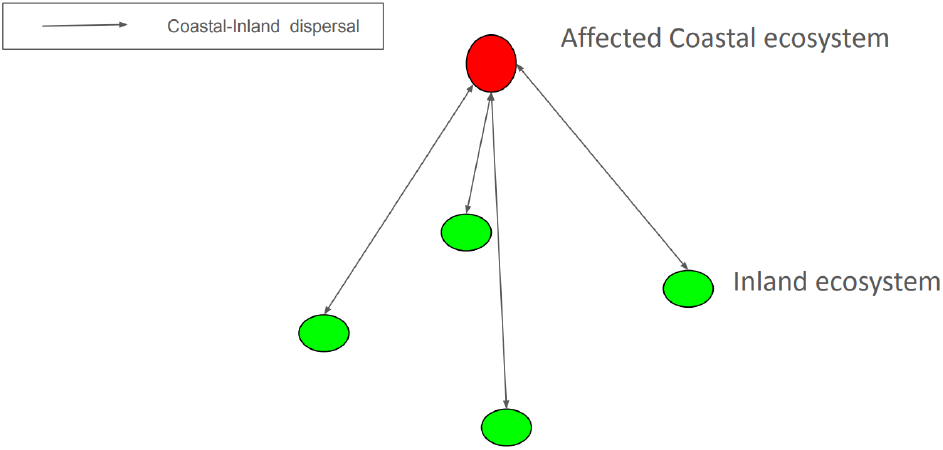
Schematic diagram representing the ecological network formed by the affected coastal ecosystem and the inland ecosystems.

The population dynamics of the *i*^*th*^ inland patch is given by Eq. (4), where *X*_0_ → *X*_*i*_, *Y*_0_ → *Y*_*i*_ and *Z*_0_ → *Z*_*i*_ and 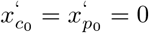. The dispersal term changes → *ε*(*Y*_0_ − *Y*_*i*_). Here, *Y*_*i*_ is the predator population of the *i*^*th*^ inland patch. Most importantly, the habitat quality of the coastal ecosystem is denoted as a function of time by *K*_0_(*t*) and for the insland ecosystem the same is given by *K*_*i*_ (constant) for the *i*^*th*^ inland patch.

### B. Modeling carrying capacity of an impacted ecosystem

The immediate impact of a tropical cyclone on a coastal ecosystem will be the mortality of a fraction of the species population. Note that the most important effect of a cyclone-originated storm surge is its impact on the habitat quality of the coastal ecosystem. Almost inevitably, the habitat quality of the coastal ecosystem is adversely affected in the long-term by the storm-surge. Cyclones create hypoxic conditions [11, 12] and storm surge that detrimentally affects the habitat quality of the coastal ecosystem [46–48] by creating either lack or redundancy of nutrients [49, 50] or aggravating the hypoxia [51, 52]. Thus, the habitat quality *K*_0_ of the ecosystem deteriorates after the impact of the storm surge. This, reasonably, long term effect on the habitat quality is modeled by a carrying capacity that deteriorates (or varies) with time. In this work, we consider three typical dynamics representing the gradually deteriorating carrying capacity. The rate of the deterioration is proportional to the storm-surge magnitude *h* and the inherent damage susceptibility of the ecosystem, quantified by the parameter *σ*, as wellthe instantaneous value of the carrying capacity. Notably, the inherent damage susceptibility is considered to be *σ* = 0 for the inland ecosystems. Thus, the deteriorating carrying capacity of the coastal ecosystem *K*_0_ is modeled as

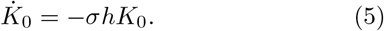

## III. RESULTS

### A. Tropical cyclone induced storm surge

Cyclones are essentially huge volumes of spiraling wind forming a vortex. A low pressure zone created in the atmosphere forms the eye of the tropical cyclone. Eventually, the wind around the eye, governed by Eq. (1), evolves into an intense, massive swirling vortex of wind with immense power. We simulate the wind-field of a cyclone, governed by Eq. (1). Figure 2 shows the cyclonic wind-field, against the backdrop of the pressure gradient given in color. For fidelity, it could be compared to wind profiles of cyclones obtained from other studies [53–56]

The intense wind-field of the cyclone creates a massive shearing stress in the atmosphere called the wind-stress.

Figure 1(a) shows pressure distribution along the wind-field of the cyclone. The thickness of the arrows, depicting the wind-flow of the cyclone, is proportional to the magnitude of the wind velocity. A very small circle at the center is the eye with relative calmness and no wind. The increasing thickness of the arrows around the center proves that the wind is strongest around the eye. The wind velocity decreases gradually from the center of the eye. This prompts that the cyclonic wind must vary radially. The radial profile of the wind-stress is shown in Fig. 1(b). The wind-stress is maximum at a particular radial distance from the eye. Close to the origin, i.e., the eye of the cyclone, the wind field is negligible, but the pressure deficit is maximum. This implies that the eye of the cyclone is relatively calm. Note that the y-axis in Fig. 1(b) has the range (0.1 − 7 × 10^7^) Pa. The ‘zero’ of the y-axis is basically the normal air-pressure. The primary physical damage caused by the tropical cyclone on making landfall is due to this wind-stress. However, since tropical cyclones are mostly created over oceans, the most important effect of this wind-stress is the formation of storm surge, i.e. elevation in sea-water levels. The dynamics of the ocean water level is governed by Eq. (3). The periodic behavior of the tide and ebb of the thermocline water level obtained from the storm-surge model when unperturbed by the extra wind-stress from the cyclone is shown green dotted line in Fig. 1(c). However, a large elevation in the height of the thermocline water level caused by the cyclonic wind-stress (refer Fig. 1 (b)) during the cyclone is evident from red solid line in Fig. 1(c). This phenomenon is called storm surge. For the particular case shown in Fig. 1(c), the storm surge corresponds to a 22 ft tidal wave.

### B. Behavior of an unaffected coastal ecosystem

First, we investigate the behavior of the coastal ecosystem when unperturbed by the extreme event. The bifurcation diagram (see Fig. 3) of the solitary, unconnected coastal ecosystem evidently display chaotic oscillation of the population density in acceptable range of the carrying capacity *K*_0_ ∈ (0.8, 0.98) when unaffected by a tropical cyclone. The dynamics saturates to a high-density steady state in the range *K*_0_ ∈ (0.99, 1.00). In this range, with increasing carrying capacity, the steady state population density keeps on increasing. For this situation, These values, unless otherwise specified are taken to be constant throughout this study. For the coastal ecosystem, the carrying capacity varies as *K*_0_ = *K*_0_)(*t*). However, at initial time at *t* = 0, we consider *K*_0_(0) = 0.97, ensuring that we start from a stable chaotic oscillatory dynamics. Figure 3 establishes that the dynamics of the unperturbed coastal ecosystem in the explored range of carrying capacity, along with the parameter values considered throughout this work, create a self-sustainable ecosystem dynamics without dispersal and are therefore within acceptable range. It is to be noted that with the carrying capacity *K*_*i*_ kept constant at *K*_*i*_ = 0.97, Fig. 3 is also representative of the dynamics of the individual inland ecosystems.

**FIG. 3.**
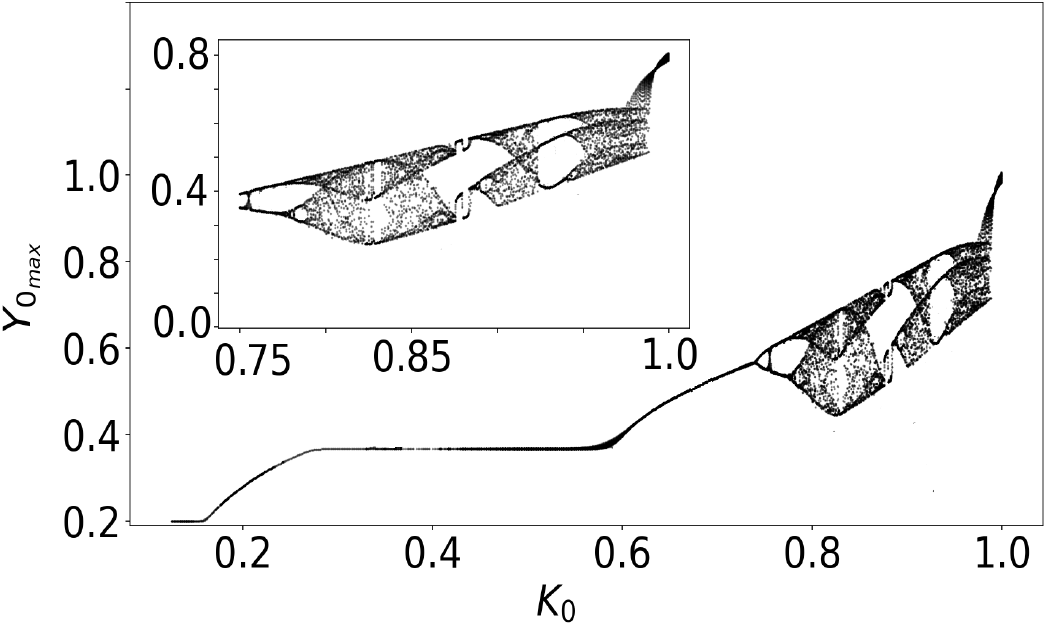
Bifurcation of an unperturbed coastal ecosystem corresponding to carrying capacity *K*_0_. The parameter values are the same as in Eqs. (4)

### C. Behaviour of two dispersally connected affected coastal ecosystem

Coastal ecosystems, close enough so that they are impacted identically by the cyclone, can be connected with each other through dispersal. We investigate if dispersal between them can also revive populations of the respective coastal ecosystem. Figure 4 demonstrates the time-series behavior of two such affected almost identical ecosystems (*X*_0_, *Y*_0_, *Z*_0_) and 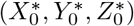, with slight variation in their damage susceptibilities (*σ* = 8 × 10^−4^ and *σ*^*^ = 7.5 × 10^−4^). From the figure, it is evident that even though considerable dispersal can boost population of both ecosystems initially, eventually, dispersal, even of considerable magnitude (*ε* = 0.2) between two affected coastal ecosystems is also not sufficient to prevent the eventual extinction of the inherent population.

**FIG. 4.**
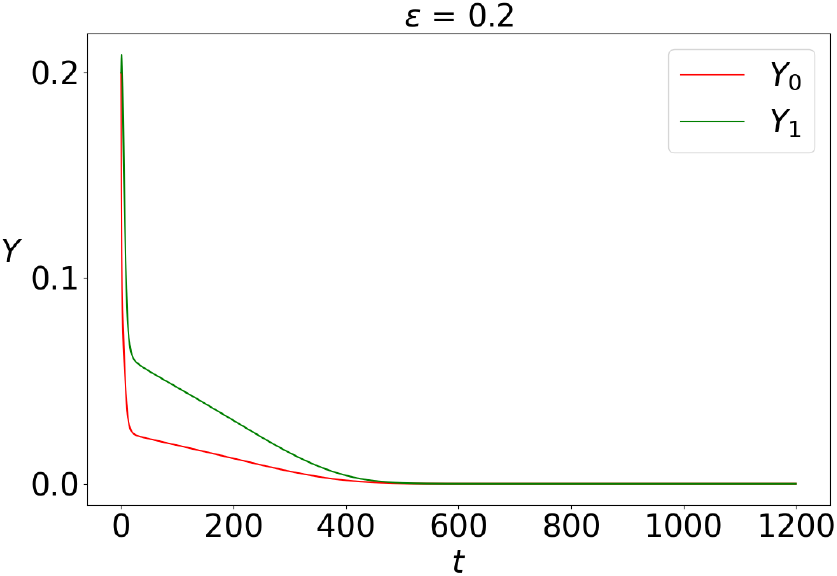
Predator population (*Y*) of two dispersally connected coastal ecosystems affected by the cyclone (extreme event). Dispersal magnitude is high *ε* = 0.2. The parameter values are the same as in Eqs. (4)

### D. Dispersal revitalizes ecosystems destroyed by tropical cyclones

In this section, we quantify the damage caused to a coastal ecosystem by a tropical cyclone and analyze how dispersal between the affected coastal ecosystem and the ‘safer’ inland ecosystems can counteract the onslaught. The food webs of the coastal ecosystem and the inland ecosystem are governed by Eq. (4).

The storm surge shown in Fig.1(c) strongly affects the habitat quality and the dynamics of coastal ecosystems in two major ways. In a more immediate time-scale, it increases the mortality of all the species living in the affected ecosystem. This is due to the immediate impact of increasing water level, salinity, or can be caused by either lack (or in rare occasions, abundance) of nutrients which is unfavorable for many coastal species. Secondly and more importantly, as a long term effect, storm surge reduces the habitat quality of the affected ecosystem by creating hypoxic conditions (as well as extended salinity and decrease of resources). Even when the conditions are not hypoxic, there can be severe damage to the habitat quality in the aftermath of storm surge. It is very important to note that this is a long-term effect, which, even if does not extend for decades, is long term enough to justify the simulation of a coastal ecosystem taking it into account ([57–60]).

In this work, we consider the behavior and revival strategies of a coastal ecosystem affected in the long term by deteriorating habitat quality. The deterioration of the habitat quality of the affected ecosystem is quantified by Eq. (5). The food web of the affected coastal ecosystem is given by Eq. (4). While the common prey (*X*_0_) of the affected coastal ecosystem food webs i.e., phytoplanktons and zooplanktons, are relatively localized, the predators (like fish) (*Y*_0_) can regularly disperse to inland ecosystems. The inland ecosystems are directly much less affected by the cyclone and are ‘safer’ [49], [61], [62]. The population dynamics of the predator (*Y*_0_) of an affected but isolated (i.e., dispersally unconnected) coastal system is shown in Fig. 5(a). Evidently, the predator population is affected extremely detrimentally with no chance of survival i.e., both amplitude and oscillation death. Thus, from a point of view of conservation of the ecosystem, the question of population revival through dispersal needs to be investigated. The predator population dynamics (*Y*_0_) when the ecosystem is a part of a metacommunity (i.e., connected by dispersal) is shown in Fig. 5(b). It is evident from the figure that in contrast to the previous figure, there is clearly a non-zero predator population density establishing that dispersal revives the predator species. The consumers (*Z*_0_) of the food web heavily depend on the predator for their survival, and indirectly so does the prey (since unchecked growth of prey is self-harming). So, along with the predator, the consequent revival of the affected ecosystem engineered by dispersal of the predators also follows.

**FIG. 5.**
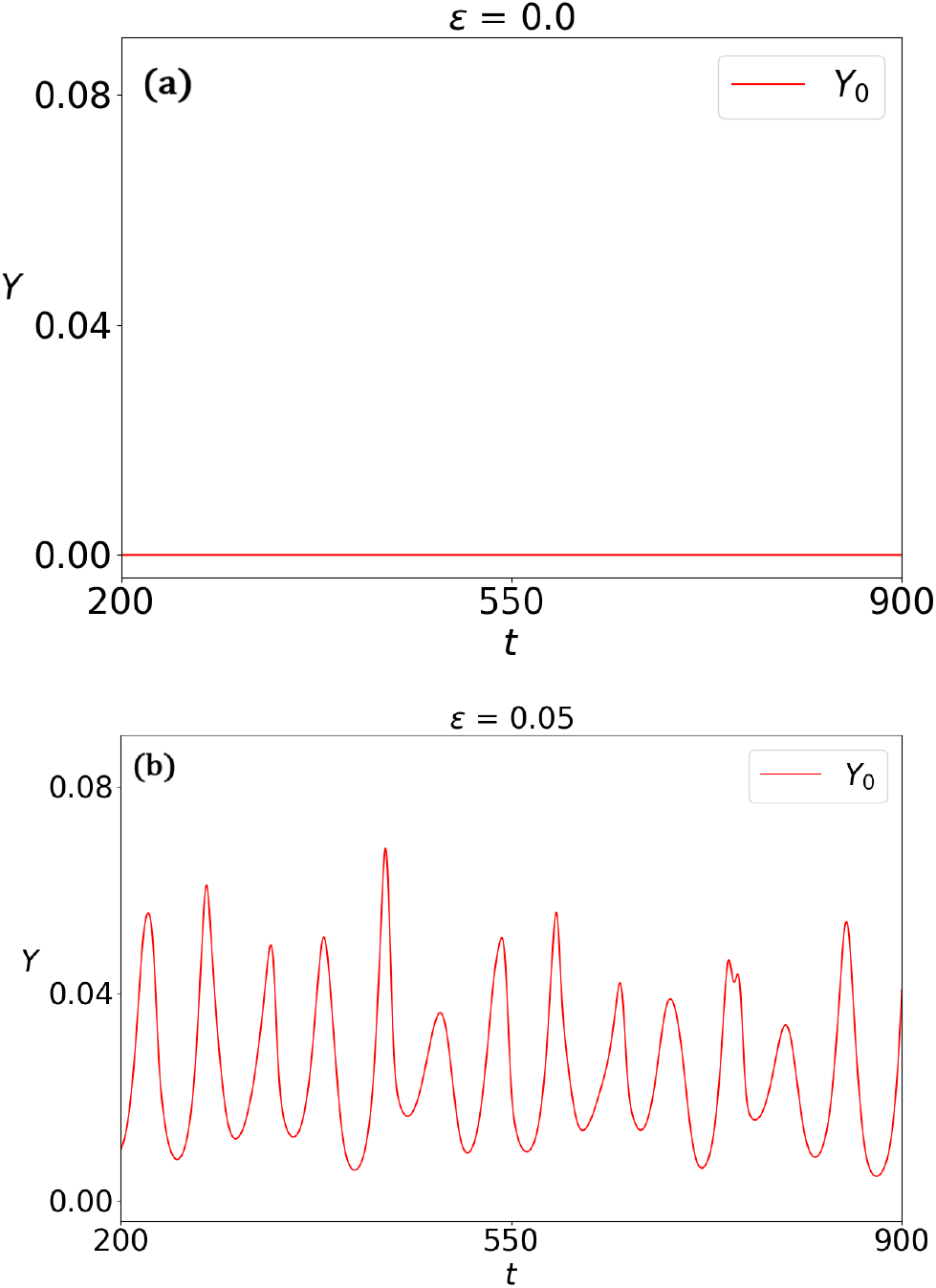
(a) Predator population (*Y*_0_) of the cyclone-affected coastal ecosystem without dispersal. (b) Predator population (*Y*_0_) of cyclone-affected ecosystem with optimal dispersal. Except for the dispersal amplitude (*ε* = 0.05), all the parameters are the same as in Eq. (4).

As the predator is revived, it has two beneficial effects on the ecosystem. Since the consumer *Z*_0_ survives due to the predator population density, non-zero predator population ensures their survival. Secondly, as the prey population *X*_0_, unchecked by predation, cannot survive due to intra-species competition, the survival of predator also ensures healthy prey population. So, the dispersal of predators ensure non-zero, healthy population of prey, predators and consumers, i.e. the whole ecosystem. Most importantly, mere non-zero dispersal does not ensure the survival of the ecosystem. The revival of the predator species onsets only when the degree of the dispersal of predators is beyond a certain threshold, which depends on numerous important factors like intensity of cyclone, intrinsic damage-proneness of the coastal ecosystem and fragmentation level of habitat, as we will demonstrate in the following.

The intensity of tropical cyclone is governed by the Holland parameter *b*_*h*_. Depending upon the intensity of cyclone, the critical dispersal value required for the revival of the predator population differs.

Low values of the Holland parameter imply weaker cyclones and correspondingly, the critical dispersal value required for species revival is small. For stronger tropical cyclones, implied by high values of the Holland parameter, larger values of optimal dispersal are necessary. The duress of extreme events makes normal levels of dispersal unattainable for the affected ecosystem. Moreover, all ecosystems may not have the same intrinsic robustness to withstand extreme events. The intrinsic damage proneness of an ecosystem is quantified by the parameter *σ* in Eq. (**??**). Depending on the degree of *σ*, the response of the coastal ecosystem to cyclones of equal intensity and consequently the critical dispersal required for the species revival also differs. Clearly, the critical dispersal value required for species revival is determined by both damage-proneness and cyclone intensity.

The exponential decrease in carrying capacity mitigates the damage to a very limited extent. Figure 6 demonstrates the degree of variation of critical dispersal corresponding to both *b*_*h*_ and *σh*. In this case, the increase in critical dispersal is governed mostly by the increase in damage susceptibility of the ecosystem. In the range of *σh ϵ* (3.5, 5.5) × 10^−4^, the critical dispersal required to revive population is of a lower degree, even though with increasing *b*_*h*_, there is a limited increase in critical dispersal level. However, For *σh >* 5.5 × 10^−4^, the required critical dispersal increases from a lower value to a higher value as a function of *σh* elucidating that a low(high) intensity cyclone along with a low(high) damage susceptibility ecosystem necessitates a low(high) degree of predator dispersal to enable the persistence of the ecosystem. However, *σhε*(6.0 → 7.5) ×10^−4^, the critical dispersal increases to a higher value (more than 3 times). Clearly, for a low intensity cyclone and a low damage-proneness, correspondingly a low critical dispersal is sufficient to counter the detrimental effects of the cyclone in the range of Holland parameter *b*_*h*_*ε*(0.1, 1.1) and *σh*(3.5 → 5.5 × 10^−4^).

**FIG. 6.**
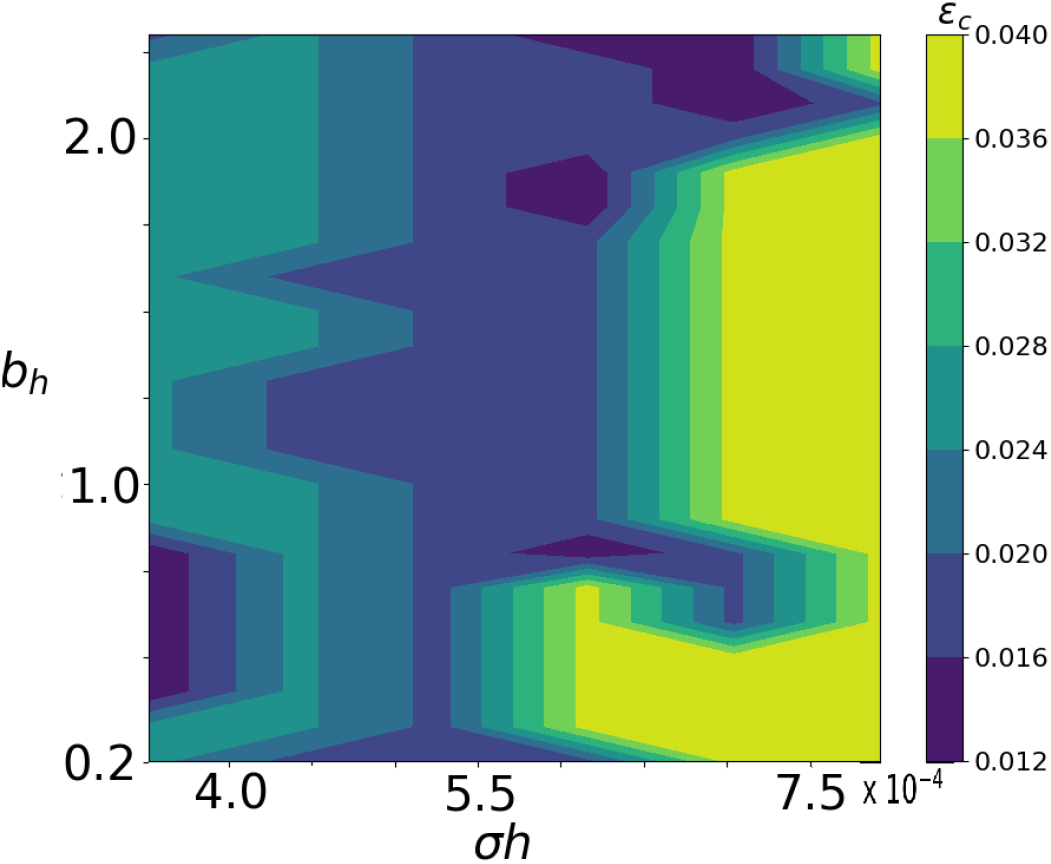
Gradual variation of the critical dispersal (*ε*_*c*_) corresponding to change in both preexisting damage-proneness of ecosystem (*σh*) and tropical cyclone intensity (Holland parameter: *b*_*h*_) when *K*_0_ follows Eq. (5). All the other parameters have values mentioned in Eq. (4).

Figure 7 demonstrates the behavior of the affected ecosystem under increasing dispersal, when *K*_0_ follows Eq. 5. Chaotic dynamics exists for the range of dispersal (*ε*0.03) explored. With increase in dispersal from *εϵ*(0.02, 0.125) however, the range of population density increases. On further increase of dispersal, the dynamics transitions from chaotic to limit cycle dynamics for the small range of *εϵ*(0.125, 0.15). From the range of *εϵ*(0.15, 0.235), the population density exhibits a chaotic dynamics and increases, with low degree of oscillation, i.e., the maxima and minima are very close. For even higher dispersal, due to the reason stated in the above paragraph, the population density of both the affected coastal ecosystem and the inland ecosystem decreases to almost comparable values for *εϵ*(0.225, 0.35). Even in this region the dynamics is chaotic. Figure 7 further demonstrates that there is a transition from chaotic to steady state dynamics for *ε >* 0.35.

**FIG. 7.**
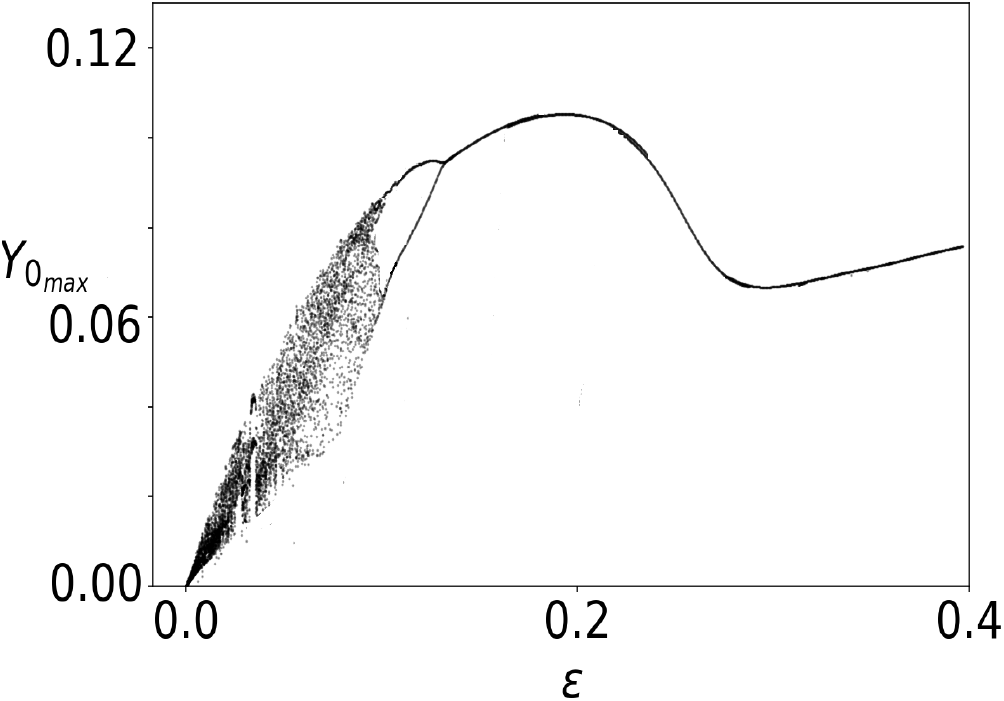
(Bifurcation diagram of the cyclone-affected coastal ecosystem corresponding to dispersal *ε* (b) when *K*_0_ follows Eq. (5). All the parameters are the same as in Eq. (4).

## IV. DISCUSSION AND CONCLUSION

We have considered the interaction of a typically complex and ecologically relevant extreme event with an affected ecosystem described by a suitable ecological model. We have examined the impact of a tropical cyclone (a climatic extreme event) on a coastal ecosystem. We establish that the coastal ecosystem has normal unperturbed behavior without the impact of the tropical cyclone. We further establish that dispersal between two affected coastal ecosystems cannot revive either of them. Next, we consider three distinct but increasingly probable dynamics followed by the deteriorating carrying capacity of the ecosystem. For all the three cases, we found that the local species of the affected coastal ecosystem suffer extinction when they remain disconnected from other ‘safer’ inland ecosystems through dispersal. Nevertheless, for all the cases we have also observed that the local population of the affected ecosystem persists above a critical dispersal rate between the former and the safer inland ecosystem. The critical dispersal rate is found to increase with the intensity of the tropical cyclone and with the increasing degree of damage-proneness of the ecosystem for a fixed intensity of tropical cyclone. For each case, we study the behavior of the affected ecosystem under increasing dispersal and establish that the ecosystem either exhibits chaotic dynamics all through (first case) or undergoes transitions from chaotic to limit cycle dynamics and finally to steady state dynamics.

It is also important to highlight another aspect of this study. Extreme events, by their definitions are of huge magnitude. For something like a cyclone, scope of its damage can only be measured after it passes. A simulation study gives us the advantage of taking precautionary measures by predicting the impact of such extreme events on ecosystems. The ecosystems that are affected by such extreme events in general are also part of spatially extended ecological networks. Moreover, the population dynamics of most ecosystems vary on a long time-scale. Observation of exact long-term impacts of the extreme events on spatially extended ecosystems through field data requires considerable effort, time and money. Added to that, such studies are often based on modeling the impact of an extreme event on a single species and may miss the increased complexity of food webs and networks. Such approaches might result in overlooking suitable conservation measures that help the total metacommunity. Hence, a simulation study such as ours, which includes the complex models of both extreme events and affected ecological networks, and also their interactions gives us a clear advantage of early predictive capability. This can provide us with much needed time to implement conservation measures in danger of an inevitable extreme event impact.

## ACKNOWLEDGMENTS

DB acknowledges IISER-TVM for infrastructural and funding support. food web of a large subtropical lake, Journal Of Plankton Research **33.7**, 1081 (2011).

